# Prolonged exposure to constant environmental conditions prompts nonrandom genetic variation

**DOI:** 10.1101/2020.04.07.030270

**Authors:** Francesco Catania, Rebecca Hagen, Valerio Vitali

## Abstract

Long-term environmental exposure under selection-free conditions has no consequences for fitness under the neo-Darwinian paradigm but it may provoke adaptive developmental buffering if environmental pressures foster directional organismal changes. To test this hypothesis, we revisited a Mutation Accumulation (MA) experiment where isogenic lines of the ciliate *Paramecium* were propagated for >40 sexual cycles (∼4 years) in a nearly selection-free and nutrient-rich environment. We find that these MA lines’ somatic genome is enriched with intervening segments of DNA (IESs), which are normally eliminated during germline-soma differentiation. Across independent replicate MA lines, an excess of these somatic IESs fall into a class of epigenetically controlled sequences, map to the same genomic locations, and preferentially disrupt loci that regulate nutrient metabolism. Although further work is needed to assess the phenotypic consequences of somatic IESs, these findings support a model where environmentally induced developmental variants may restore an adaptive fit between phenotype and environment. In this model, positive selection is surprisingly dispensable for adaptation.

## INTRODUCTION

One hundred and sixty-one years after the publication of *On the Origin of Species* **(Darwin 1859)**, the processes by which organisms adapt to their surroundings remain under scrutiny. The retention of beneficial mutations by means of natural selection is the dominant view with respect to the generation of adaptively useful variation. Against this background, a growing number of studies have proposed an additional mechanism of adaptation, which lies outside the classical neo-Darwinian scheme of allelic replacement due to selection **(West-Eberhard 2003; West-Eberhard 2005; West-Eberhard 2008)**. In this mechanism, environmental pressures expose latent developmental variation that is capable of adjusting physiology and development sensitivity to disturbances, without new genetic change. These adaptive developmental variations, when faithfully passed down across sexual generations, may become genetically assimilated or accommodated over evolutionary time **(Baldwin 1902; Schmalhausen 1949; Waddington 1953; Jablonka and Lamb 2005; West-Eberhard 2005; Pigliucci et al. 2006)**. Thus, a sufficiently long exposure to a given environmental condition (*e.g*. constant darkness) might actively mediate the emergence of phenotypes (*e.g*. the loss of eyes in cave-dwelling populations **(Stern and Crandall 2018)**), which adaptively increase energetic efficiency by restoring homeostasis **(Lotka 1922; Calow 1982; Torres 1991; Parsons 2009)**. Importantly, this mechanism makes positive selection dispensable for adaptation.

Minimal selection efficacy is the central characteristic of Mutation Accumulation (MA) experiments. In MA experiments, replicate lines with the same genotype are allowed to accumulate spontaneous mutations under conditions where each generation experiences extreme bottlenecking **(Katju and Bergthorsson 2019)**. Further, MA lines are reared in constant environmental conditions (*e.g*., high nutrient level), which may foster directional organismal changes under the environment induction-based mechanism of adaptation illustrated above. This, alongside their virtually selection-free conditions, makes MA studies arguably the best currently available tool to explore the role that the environment plays in the emergence of evolutionary innovations.

To test this idea, we revisited a published MA study where lines of the ciliate *Paramecium tetraurelia* were propagated from an isogenic state for ∼3,300 asexual generations, intercalated with >40 sexual generations, in a nutrient-rich culture medium **(Sung et al. 2012)**. The daily single-cell transfers in this study minimize the efficacy of selection, whereas the 29 putative mutations detected at the end of the ∼4-year experiment across multiple MA lines reveal particularly low levels of genetic diversity. Following each episode of sexual reproduction (self-fertilization), *P. tetraurelia* – which, like other ciliates, houses a germline and a somatic nucleus in the same cytoplasm **(Sonneborn 1957)** – gives rise to offspring that are completely homozygous and genetically identical to its parent. In *P. tetraurelia*, the development of a new somatic nucleus from the zygotic nucleus involves DNA amplification (from 2*n* to ∼860*n* **(Woodard et al. 1961)**) and rearrangements including the elimination of ∼45,000 intervening germline DNA regions termed Internal Eliminated Sequences (IESs) **(Betermier 2004; Duharcourt et al. 2009; Arnaiz et al. 2012)**. This process of Programmed DNA Elimination (PDE) is not foolproof, allowing newly developed somatic genomes to retain IESs **(Duret et al. 2008; Catania et al. 2013; Vitali et al. 2019)**. In a typical deep-sequencing study, IES retention is limited to a few hundred somatic loci and mainly involves ≤ 5% of the ∼860 copies *per* locus when *P. tetraurelia* is cultured under standard conditions (**Table S1**). However, the cultivation environment can significantly perturb PDE in *P. tetraurelia*, generating DNA variation that might be heritable and potentially adaptive **(Vitali et al. 2019)**.

## RESULTS

### An excess of IESs accrue in the somatic nucleus of the MA lines

We asked whether the long-term exposure of *Paramecium* to the MA regime is coupled with an enhanced accrual of IESs in the somatic genome. After filtering out somatic loci with a trivial fraction of IES-retaining mapping reads (arbitrarily set to ≤5% or with IES retention Score [IRS] ≤ 0.05), we found that the surveyed MA lines exhibit between 2 and 4 times more IES-containing loci than the control *P. tetraurelia* stocks, one of which is d4-2, the same stock used for the MA experiment (**Figure 1A**). This enrichment is largely confined to IESs with 0.05 < IRS ≤ 0.2 (two proportions Z test, *P* < 0.0001; **Figure 1B**). Further, we detected differences in the magnitude of IES retention at tens of loci. IES retention can either increase (**Figure 1C**) or decrease (**Figure 1D**) significantly in the MA lines compared to the control stocks as well as among the control stocks (**Figure 1C, 1D**). These findings suggest that the somatic nucleus of *P. tetraurelia* undergoes recurrent and pronounced seesaw IES retention dynamics, extending previous across-species observations **(Catania et al. 2013)**.

**Figure 1.**
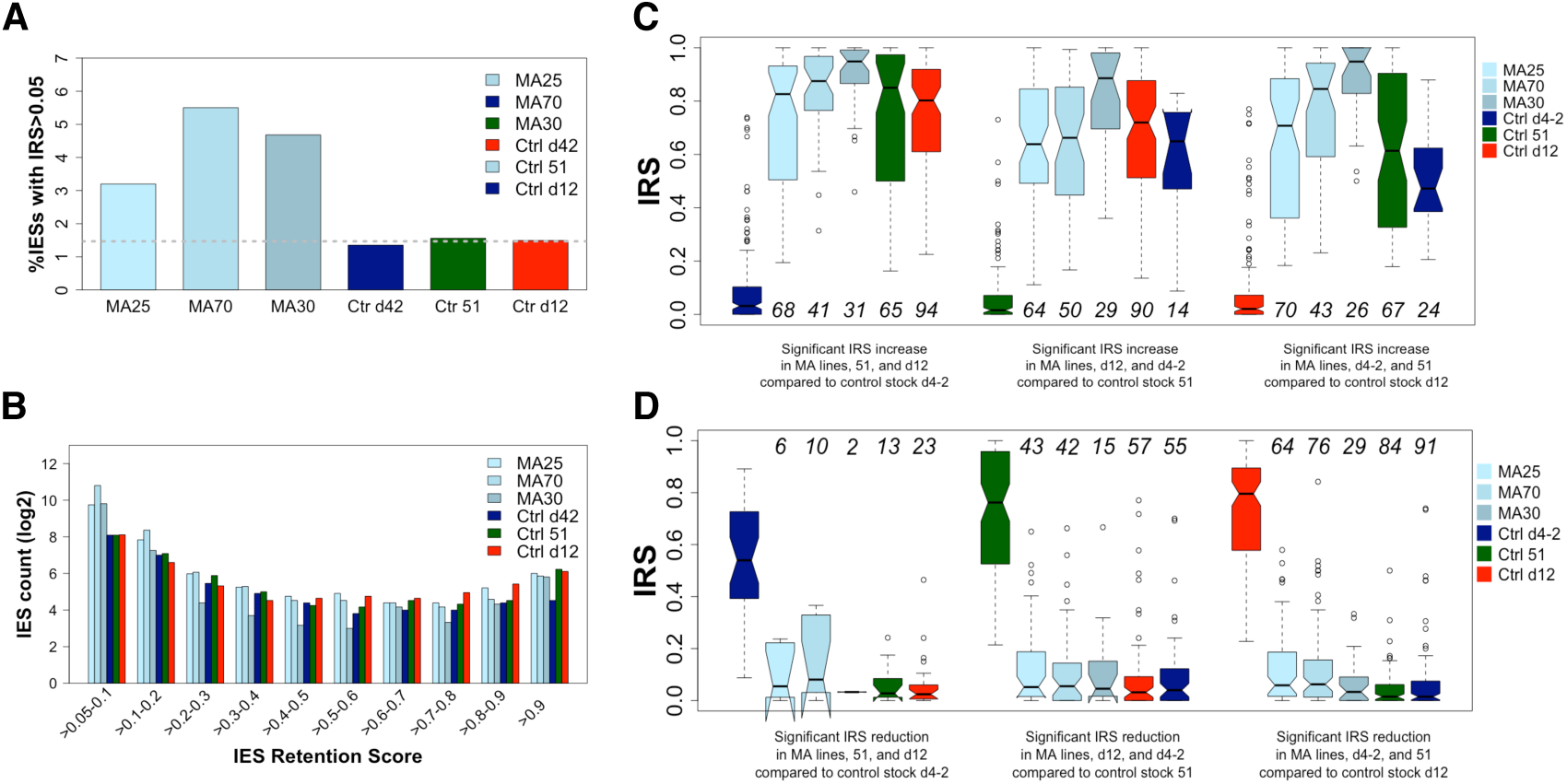
MA lines contain a higher percentage of somatic IESs with IES Retention Score (IRS) > 0.05 compared to ∼isogenic control *P. tetraurelia* stocks d4-2 (the same used as progenitor of the MA lines) 51 and d12 **(A)**. MA lines display a relatively higher count (log2) of somatic IESs with 0.05<IRS≤0.2 **(B)**. At ten of loci (numbers in italics), the IES Retention Scores in the MA lines are significantly elevated (**C**) or significantly reduced (**D**) compared to control stocks.

### Developmental changes recur across replicate lines

We also found that the somatic IESs in the MA lines greatly overlap (**Figure 2**). When simulating IES retention as a fully stochastic process, we estimated that independent replicate MA lines retain in their somatic nucleus more of the same IESs than would be expected by chance (Z-score; *P*<0.0001). Similarly, an excess of overlapping IESs among the MA lines is observed for IESs with significantly enhanced or significantly reduced levels of retention in the MA lines compared to the parental stock d4-2 (**Figure S1**).

**Figure 2.**
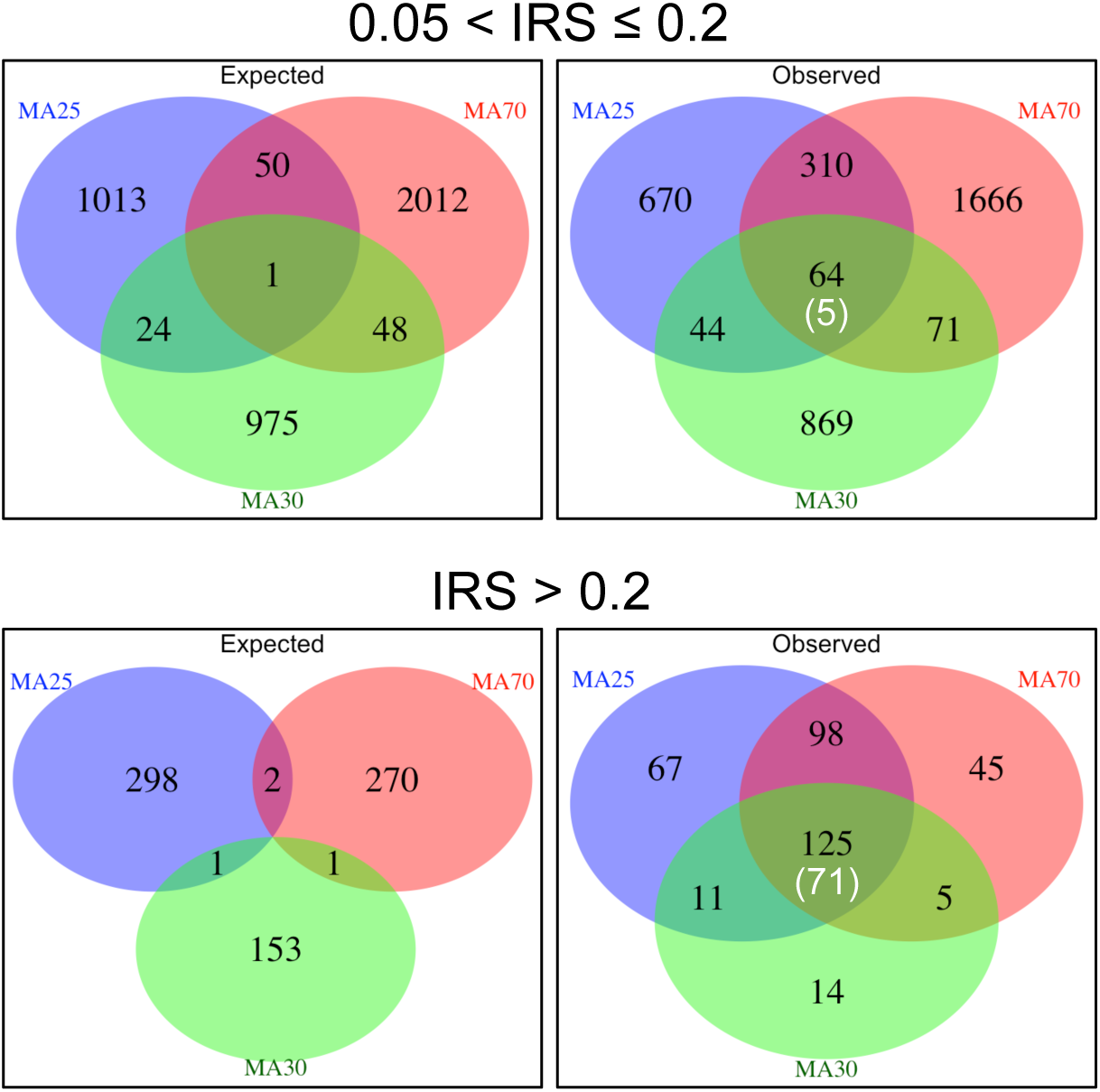
Intersections between somatic IESs in the MA lines with IES Retention Score (IRS) ranging between >0.05 and 0.2 and >0.2. Expected values were calculated from 10,000 simulations. Only IESs supported by >20 reads were considered. The number of 6-way shared IESs (*i.e*., including the MA lines and the control stocks d4-2, 51, and d12) is within parentheses.

Additional observations strengthen the hypothesis that somatic IESs with 0.05 < IRS ≤ 0.2 have accrued non-randomly or have undergone non-random dynamics in the MA lines. First, of the 64 IESs with 0.05 < IRS ≤ 0.2 that recur in the somatic nucleus of the MA lines, 23 (36%) have comparable retention scores in the parental stock versus 85 (68%) for recurring IESs with IRS > 0.2. Second, 23% of the IESs with 0.05 < IRS ≤ 0.2 in the MA lines are never retained in the parental stock despite high coverage (≥53 reads, on average) versus 4% of the IESs with IRS > 0.2. Last, somatic IESs that are shared among MA lines and with 0.05 < IRS ≤ 0.2 are less likely to be retained in all the three control stocks to a similar extent compared to IESs with IRS>0.2 (8% vs. 57%).

Altogether these observations suggest (*i*) that numerous developmental changes accumulated in the MA lines in parallel during the course of the ∼4-year long experiment, and (*ii*) that it is unlikely that all of the detected incomplete IES excisions are mere errors.

### Recurrent somatic IESs in the MA lines are not the product of weak *cis*-acting IES recognition/excision signals

The observed non-randomness of incomplete IES excisions in the MA lines (class 0.05 < IRS ≤ 0.2, in particular) is puzzling. Parallel changes are often considered to result from selection and adaptation **(Losos 2011)**. But how can the same developmental changes across independent replicate lines be the result of selection in an experiment where selection efficacy is severely diminished? One alternative explanation is that IESs that are retained in the somatic nucleus of the MA lines have particularly weak *cis*-acting DNA splicing signals, which favor their reappearance at every sexual generation. Our observations do not align well with this possibility: IESs with 0.05 < IRS ≤ 0.2 exhibit stronger (rather than weaker) *cis*-acting excision/recognition signals in the MA lines compared to the control stocks (C_in_-scores **(Ferro et al. 2015)**: 0.599 vs. 0.556, t-test, *P* < 0.0001). The estimated signal strength is comparable for IESs with IRS > 0.2 (C_in_-scores: 0.506 vs. 0.504, t-test, *P* = 0.878).

### Somatic IESs in the MA lines map preferentially within coding exons and may impact gene expression levels

Somatic IESs that recur across the MA lines might reflect adaptation to the MA regime. If so, then these IESs are likely to affect the phenotype. Although the non-availability of the MA lines prevents us from testing this expectation directly, indirect observations are consistent with this. For one, we found that the somatic IESs that are over-represented in the MA lines (0.05 < IRS ≤ 0.2) reside most frequently within coding exons (**Figure 3A**), in striking contrast to the control stocks where somatic IESs with comparable IRSs are most frequently located in intergenic regions (χ^2^= 181.7, df = 2, *P*<0.0001; **Figure 3B**). The detected positional bias in the MA lines is associated with the introduction of premature termination codons (PTCs) in the nascent transcript: 77% of the ORFs acquire PTCs upon IES retention (**Figure 3C**). This suggests that many IES-containing transcripts in the MA lines are targeted for degradation by the cellular surveillance systems. It also predicts that the MA lines and the control stocks may have undergone divergent changes in gene expression level over the ∼4-year long MA experiment. To test this, we leveraged the transcriptomes of the control stocks 51 **(Arnaiz et al. 2017)** and d12 **(Vitali et al. 2019)** obtained at standard (MA experiment-like) cultivation conditions. We found that genes with reduced levels of expression in the control stocks preferentially accumulate somatic IESs (**Figure 3D**), consistent with previous reports **(Arnaiz et al. 2012; Ferro et al. 2015; Vitali et al. 2019)**. The opposite is true for the MA lines, where an excess of highly expressed genes retain IESs relative to the control stocks (two proportions Z test, *P* < 0.0001). Further, the amount of somatic IESs in MA line genes decreases gradually with gene expression level, rather than increasing as in the control stocks (**Figure 3D**). These results suggest that IES retention may have re-configured part of the transcriptional landscape in the MA lines. Alternatively, somatic IESs may have accrued preferentially in down-regulated genes. At any rate, the negative scaling between IES retention levels and gene expression in both the control stock d12 **(Vitali et al. 2019)** and stock 51 (Kendall’s τ = −0.176, *P*<0.0001) further supports the relationship between developmental variation and somatic gene expression in *P. tetraurelia*.

**Figure 3.**
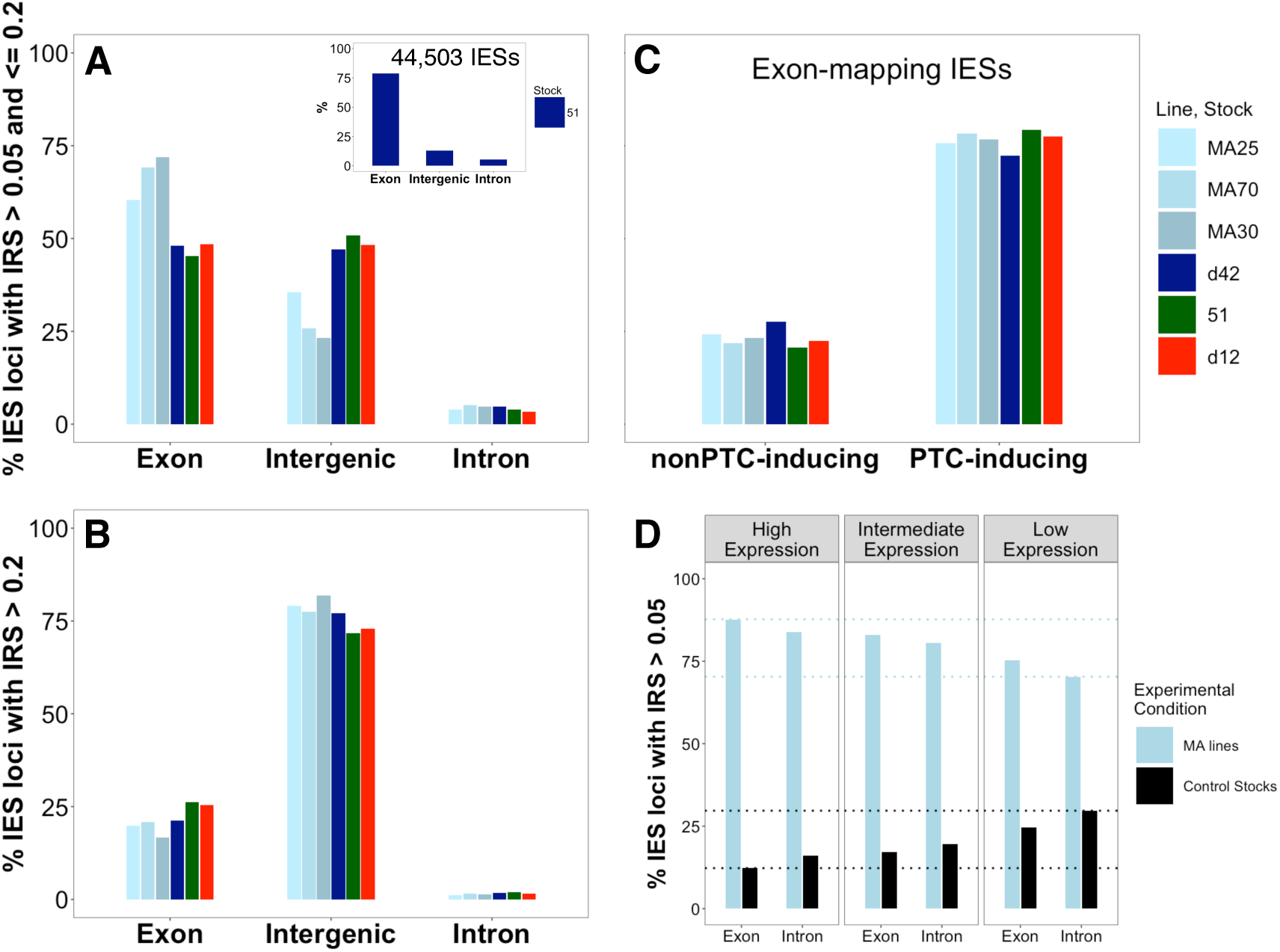
Genomic distribution of somatic IESs with IES Retention Score (IRS) ranging between >0.05 and ≤0.2 (**A**) and >0.2 (**B**) in MA lines and control stocks. The vast majority of exon-mapping somatic IESs with IRS >0.05 and ≤0.2 disrupt the ORF (**C**). Relationship between level of gene expression (as in stocks 51 and d12) and percentage of somatic IESs (IRS > 0.05). Highly expressed genes accumulate fewer incompletely excised IESs compared to weakly expressed genes in the stocks d4-2, 51, and d12. The reverse trend is detected for the MA lines (average values are shown) (**D**).

### The retention of gene-adjacent intergenic IESs may alter regulatory sequences

Similar to intragenic IESs, somatic IESs that map to gene-flanking intergenic regions might also alter gene expression level in the MA lines. The positive scaling between gene expression level and IES distance to gene 5’ end (**Figure 4A**) aligns with this possibility, and so does the negative relationship between the IRS of intergenic IESs and the expression levels of genes with the 5’ end facing the IES (**Figure 4C**). More explicitly, genes that reside downstream from somatic IESs appear to experience a more pronounced down-regulation when the IES retention score is more elevated. Notably, none of the relationships described above is statistically significant downstream from the gene 3’ end, where promoters do not occur (**Figure 4B, 4D**). These and the preceding findings suggest that both intragenic and intergenic IESs, when retained, are likely to weaken the expression of a non-random set of proteins. We postulate that these expression changes contribute to a new phenotypic state.

**Figure 4.**
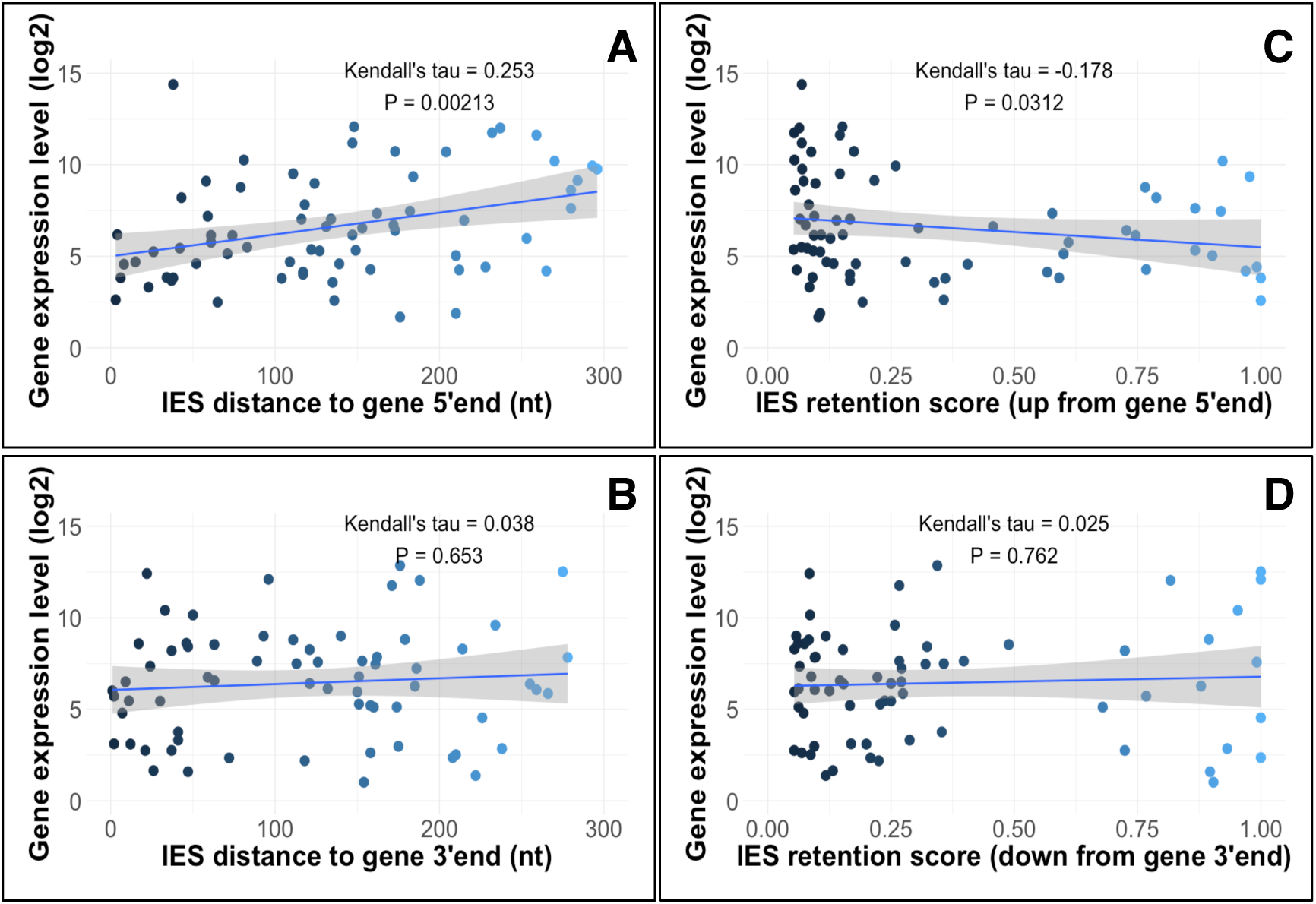
Relationship between gene expression levels (stock 51, vegetative stage) and *i*) IES distance from the corresponding gene 5’ end (**A**) and 3’ end (**B**) or *ii*) IES Retention Score (0.05 < IRS ≤ 1) within 300nt from the gene 5’ end (**C**) or 3’ end (**D**).

### Developmental changes in the MA lines target genes that may restore metabolic homeostasis

We next determined the cellular functions that IES retention is likely to impact (**Table S2**). Concerning the intragenic somatic IESs with 0.05 < IRS ≤ 0.2, we detected an MA line-exclusive association with cytoskeleton and membrane components as well as microtubule-based movement and organic substance biosynthetic process. Molecular functions such as endopeptidase inhibitor activity (GO:0004866); serine-type endopeptidase inhibitor activity (GO:0004867); and peptidase inhibitor activity (GO:0030414) are enriched in the parental stock d4-2 but are missing in the MA lines. Further, an epidermal growth factor (EGF)-like domain is enriched among the MA line genes that bear intragenic IESs with 0.05 < IRS ≤ 0.2 (Fisher’s exact test, *P*<0.0001), but is neither detected for the MA line genes that contain IESs with IRS >0.2 nor for the IES-retaining genes of the parental stock d4-2 or the control stock 51. This latter observation is interesting because previous reports suggest that *P. tetraurelia* secretes a factor with mitogenic effects **(Tanabe et al. 1990)** that may resemble the mammalian EGF **(Tokusumi et al. 1996)** (in mammals, EGF can accelerate cell proliferation, cell migration, and may play critical roles in oocyte maturation, fertilization and embryo development). Concerning the somatic IESs in intergenic regions (≤300nt upstream from the 5’ end), in the MA lines they reside preferentially upstream of genes encoding factors that are involved in carbohydrate derivative, ATP, and ribonucleotide binding as well as integral components of the peroxisomal membrane, with major roles in lipid biosynthesis and fatty acid oxidation. Overall, these observations raise the possibility that the nutrient metabolism (*e.g*., nutrient intake, breakdown, conversion of nutrients into energy) is attenuated in the MA lines.

To gain further insight into the cellular functions that IES retention is likely to alter, we zeroed in on IESs whose levels of somatic retention between the MA lines and the parental stock d4-2 were divergent. We found two IESs that are under-represented in the somatic genome of all the MA lines relative to the parental stock (**Figure S2**). The only intragenic IES maps to a *P. tetraurelia* gene with currently unknown function (*PTET.51.1.G1120131*). This single-copy gene contains a predicted trans-membrane helix that spans the differentially retained 45nt long exon-mapping IES. The pronounced IES retention in the parental stock is predicted to perturb the topology of the encoded trans-membrane protein (**Figure S3**). It follows that *PTET.51.1.G1120131* in the MA lines might generate relatively larger amounts of a trans-membrane protein whose function is unknown. The other (intergenic) IES flanks a gene (*PTET.51.1.G1590064*) also with unknown function, which has a predicted trans-membrane domain (**Table S3**).

Eighteen IESs are fully or largely absent from the somatic DNA of the parental stock but have high levels of somatic retention in all the MA lines (**Figure S4**). Thirteen IESs are intergenic and most often flank gene 3’ ends with unknown implications (**Table S4**). The remaining five IESs interrupt five genes, three with a putative description (**Figure S4**): 1) *PTET.51.1.G0280259* is single-copy and encodes a WD repeat-containing protein that is involved in ribosome biogenesis in yeast; 2) *PTET.51.1.G0310293* encodes a guanine nucleotide exchange factor that is involved in signaling pathways such as cell proliferation; and last, 3) *PTET.51.1.G0480149* is a *CYClin* PHO80-like gene that is highly conserved from unicellular to multicellular species.

*PTET.51.1.G0480149* is interrupted in the MA lines but neither in the parental stock nor in the other control stocks (**Figure S4**). Interestingly, the cyclin PHO80-like domain (IPR013922) breaks down upon the retention of the 76nt-long IES **(Arnaiz and Sperling 2011)**. In other eukaryotes, the gene PHO80 plays a role in development; it is a key effector of the so-called PHO pathway — by which phosphate availability is sensed — and a positive regulator of the insulin-signaling pathway. This gene has three presumably functional ohnologs in *P. tetraurelia* (stock 51). Thus, the disruption of *PTET.51.1.G0480149* in the MA lines may contribute to reducing — rather than fully abolishing — *CYClin*’ s activity, possibly blunting nutrient signaling sensors and slowing development down. Overall, these observations support the idea that parallel germline DNA insertions in the somatic genome of the MA lines help mitigate their energy metabolism and growth.

### An excess of somatic IESs in the MA lines fall into a class of epigenetically controlled IESs

Finally, we asked how extensively IESs that accumulated in the MA lines’ somatic genome overlap with IESs that are epigenetically controlled in the control stock 51 **(Lhuillier-Akakpo et al. 2014; Sandoval et al. 2014; Maliszewska-Olejniczak et al. 2015)**. We found that a significant excess of the IESs that are retained in the MA lines fall into a class of IESs that are under the control of small RNAs (epi-IESs) in stock 51 (**Figure S5**). This enrichment of epi-IESs is also detected for MA lines’ somatic IESs with greatly increased (but not with greatly reduced) IRS (**Figure S5**). Previous studies have shown that epigenetically controlled somatic IESs in *P. tetraurelia* may be inherited **(Duharcourt et al. 1995; Duharcourt et al. 1998)**.

## Discussion

In an experimental setting where the efficacy of natural selection is maximally minimized **(Katju and Bergthorsson 2019)** and new genetic variation is negligible **(Sung et al. 2012)**, we found that independent replicate lines of *P. tetraurelia* accrue an excess of the same developmental variants during the course of a ∼4-year MA experiment. Such developmental variants fall into a class of sequences that may be trans-generationally inherited. Further, these variants hit preferentially somatic genes whose expression modulation might improve the match between >40 sexual generation-old *Paramecium* lines and the nutrient-rich and low population density environment of the MA experiment.

It is difficult to leverage the classic new-Darwinian account to explain these observations. On one hand, the reduced efficacy of selection in the MA experiment might have favoured the accumulation of IESs in the somatic nucleus of the MA lines. Even so, however, it remains unclear why the accrued developmental variants *i*) have relatively strong *cis*-acting DNA splicing signals, *ii*) fall largely into a class of epigenetically regulated sequences, and *iii*) are associated with, and most often appear to down-regulate, non-random cellular functions.

On the other hand, our observations are compatible with a model where habitual environmental exposure can induce molecular changes that advantageously restore metabolic homeostasis, in the near absence of selection. Future studies will have to further substantiate this model, test its generalizability and elucidate its connections with a neo-Darwinian account. It is possible that a mechanism of adaptation via environmental induction evolved via a traditional scheme of mutation and positive selection. In particular, mutation and natural selection might have facilitated the origin of an epigenetically controlled developmental program, which persists in modern paramecia and is induced by and favors adaptation to new conditions. The operationalization of this adaptive developmental program would not require positive Darwinian selection.

Without relinquishing the central role of mutation and positive selection in evolution, our study calls for a better appreciation for the significance of habitual environmental exposure at the onset of adaptations. The mechanism of adaptation that our observations point to could help explain the prevalence of rapid adaptive responses and convergent evolution in nature. It could also foster greater awareness of (and develop innovative measures against) the long-term effects of environmental changes.

## Methods

### *Paramecium* stocks

Details concerning the culture conditions, the macronuclear DNA isolation and the whole-genome sequencing of the Mutation Accumulation (MA) *Paramecium tetraurelia* lines were previously reported **(Sung et al. 2012)**. Three *Paramecium tetraurelia* MA lines were selected for this study based on their relatively higher median IES coverage (MA25, MA70, MA30; **Table S1**). The control, un-evolved, *Paramecium tetraurelia* stocks d4-2, 51, and d12 were used as control. Stock d4-2 (the same stock as the MA lines) and stock d12 are derivatives of stock 51 **(Sonneborn 1974; Rudman et al. 1991)**.

### IES datasets

The datasets of Internal Eliminated Sequences (IESs) for the control stocks were obtained from **(Arnaiz et al. 2012)** (stock d4-2), **(Lhuillier-Akakpo et al. 2014)** (stock 51, cultured at 27°C) and **(Vitali et al. 2019)** (stock d12, F0 line cultured at 25°C). The IES datasets for the MA lines were extracted with ParTIES **(Denby Wilkes et al. 2016)**. ParTIES was also used for the estimation of IES Retention Scores (IRSs), *i.e*., the *per*-locus ratio between IES-containing reads and the total number of mapping reads. IES loci with significantly different retention levels in an MA line compared to a control stock or between control stocks were designated as described in **(Vitali et al. 2019)** using a 95% confidence interval around the IRS value of the control stock. IES loci supported by <20 sequence reads were excluded. Only IESs with a size larger than 25 nucleotides are considered for this study.

### Parallel IES retention profiles

The expected number of incompletely excised IESs that occupy the same position in the somatic genome of three independently cultured *P. tetraurelia* lines was calculated by simulating IES retention over a single rearrangement as a fully stochastic process with R (version 3.2.1**(2018)**). For each of the surveyed MA lines, 10,000 random samples with size equal to the observed number of incompletely excised IESs with IRS>0.05 were extracted without replacement from the whole known set of ∼45,000 IESs in *P. tetraurelia* **(Arnaiz et al. 2012)**. The mean and the standard deviation of the distribution of IES numbers that are shared by chance across the three MA lines were used to calculate the probability of observing the actual three-way shared IES number.

### Gene expression data and GO-term enrichment analysis

Gene expression data for stocks 51 and d12 were obtained from previous studies **(Arnaiz et al. 2017; Vitali et al. 2019)**. The functional enrichment analyses of somatic-IES containing genes or genes with their 5’ end mapping ≤300nt downstream from intergenic IESs was performed using the functional annotation tools PANTHER **(Mi et al. 2017; Mi et al. 2019)** and DAVID **(Huang da et al. 2009)**. In all cases, Fisher’s exact test estimates the over-representation of GO-terms. The reference gene list associated with the set of intragenic somatic IESs is limited to set of IES-genes in *P. tetraurelia* stock 51 as IES retention cannot occur in IES-free genes (we previously demonstrated that IES-genes are enriched in specific cellular functions **(Vitali et al. 2019)**). The reference gene list associated with the set of intergenic somatic IESs includes all the genes in *P. tetraurelia* stock 51.

## Acknowledgments

This work was funded by the Deutsche Forschungsgemeinschaft (DFG, German Science Foundation) – 281125614/GRK 2220.

## Author contributions

F.C. conceived the study and carried out computational analysis, supervised, wrote the manuscript, and secured funding. R.H. and V.V. performed the data acquisition and wrote the manuscript.

